# An epitope-enriched immunogen increases site targeting in germinal centers

**DOI:** 10.1101/2022.12.01.518697

**Authors:** Timothy M. Caradonna, Ian W. Windsor, Anne A. Roffler, Shengli Song, Akiko Watanabe, Garnett Kelsoe, Masayuki Kuraoka, Aaron G. Schmidt

## Abstract

Antibody immunodominance is the asymmetric elicitation of responses against protein antigens. For influenza hemagglutinin (HA), antibody responses often target variable regions on HA and do not provide lasting protection. Next-generation influenza vaccines should elicit antibodies targeting conserved regions such as the receptor binding site (RBS). Understanding how presenting an epitope on a rationally-designed immunogen influences immune responses could help achieve this goal. Here, we compared an engineered RBS-enriched immunogen and its non-enriched counterparts to characterize RBS-directed responses. We found that enriching the RBS-epitope on a single immunogen preferentially expands RBS-directed responses relative to a cocktail of the non-epitope-enriched immunogens. Single B cell analyses showed a genetically diverse RBS-directed population that structural characterization showed engagement of the RBS with canonical features shared with both its receptor and human broadly neutralizing antibodies. These data show how epitope-enriched immunogens can expand responses to a conserved viral site, while maintaining genetic and structural diversity.

## INTRODUCTION

Immunodominance refers to the preferential elicitation of antibody responses against particular epitopes. Epitopes that are more frequently targeted by antibodies are referred to as immunodominant, while those that form minor portions of the overall repertoire are considered subdominant. For influenza virus, this phenomenon is a contributing factor to the suboptimal efficacy of current seasonal vaccines; elicited antibodies primarily target highly variable influenza hemagglutinin (HA) epitopes, which can lead to viral escape (Angeletti et al., 2017; Koel et al., 2013; Kuraoka et al., 2016; Wrammert et al., 2008). Conserved regions on HA, such as the receptor binding site (RBS) and stem, play key roles in mediating viral entry. While antibodies targeting these sites can be broadly protective, they often are a minority of the repertoire, particularly those recognizing the stem region (Andrews et al., 2015; Corti et al., 2011; Kallewaard et al., 2016; McCarthy et al., 2018; Schmidt et al., 2015b). A goal for next-generation influenza vaccines is to design immunogens that alter patterns of immunodominance to favor antibody responses against conserved epitopes.

Current immunogen design approaches include modifying HA to create minimal (e.g., stem- or “head-only”), chimeric, or hyperglycosylated constructs that preclude the *de novo* elicitation or memory recall of antibody responses to undesired, variable epitopes (Bajic et al., 2019; Corbett et al., 2019; Hai et al., 2012; Yassine et al., 2015). These approaches block B cells targeting variable epitopes from entering the germinal center (GC) that would otherwise outcompete B cells targeting conserved, protective epitopes. Alternative strategies confer an advantage to desired B cells rather than removing potential competitors; so called ‘germline-targeting’ immunogens for HIV Env, for example, expand specific B cell populations, but this approach has not been demonstrated for influenza HA (Corbett et al., 2019; Jardine et al., 2013; Jardine et al., 2016; Jardine et al., 2015). However, while germline-encoded responses are elicited by immunization, these populations often remain subdominant and their selective expansion can lead to a genetically restricted response (Harshbarger et al., 2021; Sangesland et al., 2019; Sui et al., 2009).

Complementary design strategies to elicit cross-reactive B cells, used multimeric display of antigenically distinct HA head- or stem-only constructs on a nanoparticle to produce a single immunogen (Boyoglu-Barnum et al., 2021; Kanekiyo et al., 2019). For head-only HA constructs, the predominant response targeted the ‘lateral patch’ epitope, a conserved region across the H1 subtype (Kanekiyo et al., 2019; Raymond et al., 2018). A possible interpretation of this observation is that the antigenically distinct displayed HAs created a “gradient” of epitope conservation ranging from unique to fully conserved. In other words, epitopes like the lateral patch are shared, while other variable head epitopes are not: cross-reactive B cells recognizing the former were therefore selectively enriched; despite the RBS being conserved, this approach failed to selectively enrich responses to the conserved RBS.

We previously developed “resurfaced” HA (rsHA) immunogens where antigenically distinct, non-circulating avian HAs served as scaffolds to present the conserved H1 RBS epitope (Bajic et al., 2020). While immunization yielded increased breadth of RBS-directed antibodies, we observed no increase in their overall frequency. We therefore expanded on this resurfacing approach with the goal of increasing the overall frequency of RBS-directed responses. To that end, we designed an “epitope-enriched” immunogen called a “rsHAtCh” or a resurfaced HA trimeric chimera; this immunogen contains three antigenically distinct rsHAs each presenting the same H1 RBS epitope (Caradonna et al., 2022). We hypothesized that enriching a single, conserved epitope on an immunogen would preferentially expand B cells targeting this epitope. In a humanized IGHV1-2 HC2 knock-in mouse with pre-existing immunity to H1 influenza, boosting with rsHAtCh expanded RBS-directed antibodies in both the H1-reactive serum and B cell compartments, however, the elicited responses had limited breadth (Caradonna et al., 2022). As one immunogen increased the breadth of RBS-directed responses, while the other increased the frequency, a more complete understanding is necessary to identify how best to accomplish both.

Here, we immunized human J_H_6 knock-in (KI) mice with rsHAtCh or a cocktail of its individual components as homotrimers and then characterized GC responses to understand how antibody repertoires changed. Analysis of primary GC responses show that the rsHAtCh expands RBS-directed responses relative to the rsHA homotrimer cocktail. Furthermore, rsHAtCh-elicited primary and secondary GC responses were genetically diverse and contained distinct clonally related RBS-directed B cells with sequence features of human RBS-directed broadly neutralizing antibodies (bnAbs) including engaging the RBS with receptor-like contacts. While RBS-directed antibodies isolated from primary GCs had limited breadth, those from secondary GCs had high affinity for historical H1s spanning twenty years. Selectively presenting multiple copies of a conserved epitope on a single immunogen with heterologous scaffolds expand antibody responses with enhanced breadth. These data suggest that epitope-enriched immunogens may be used to influence immune responses that favor expansion of antibody responses to conserved viral regions.

## RESULTS

### Cocktail and rsHAtCh immunizations

To understand how an epitope-enriched immunogen may influence antibody responses within a GC, we immunized three cohorts of huJ_H_6 KI mice (**Figure 1A-B**). The huJ_H_6 segment encodes a string of tyrosines in the HCDR3 and is a signature of human RBS-directed human antibodies (McCarthy et al., 2018; Schmidt et al., 2015b); we previously used these KI to characterize our first-generation homotrimeric rsHAs (Bajic et al., 2020). The first cohort received an equimolar cocktail of antigenically distinct H3, H4, and H14 HA heads “resurfaced” with the H1 RBS and cystine-stapled into homotrimers. The second cohort received an equivalent amount of our epitope enriched immunogen: rsHAtCh (Caradonna et al., 2022). Importantly, the individual components of the heterotrimer are identical to the rsHAs used in the first cohort, therefore the same overall epitopes are present in each immunization regimen. Thus, we reasoned that any observed differences in immunogenicity and changes in elicited responses between these two cohorts would be a consequence of how the components were arranged: the epitopes will either be presented together simultaneously (rsHAtCh, “chimera”) or separately (rsHA “cocktail”) within a GC. The third cohort received a homologous rsHAtCh prime-boost to test the effects of repeated exposure to an epitope-enriched immunogen. GC B cells were isolated from draining lymph nodes and single cell sorted into Nojima cultures at 8, 15, or 64 days after immunization to characterize the primary and secondary GC B cell populations, respectively (**Figure 1C**). Cocktail and chimera immunized mice had similar total GC B cells: ∼2.5×10^5^ and 7.5×10^4^ on day 8, 15, respectively. Day 64 GCs, however, had ∼9×10^3^ GC B cells, a statistically significant decrease relative to both primary GC timepoints.

**Figure 1:**
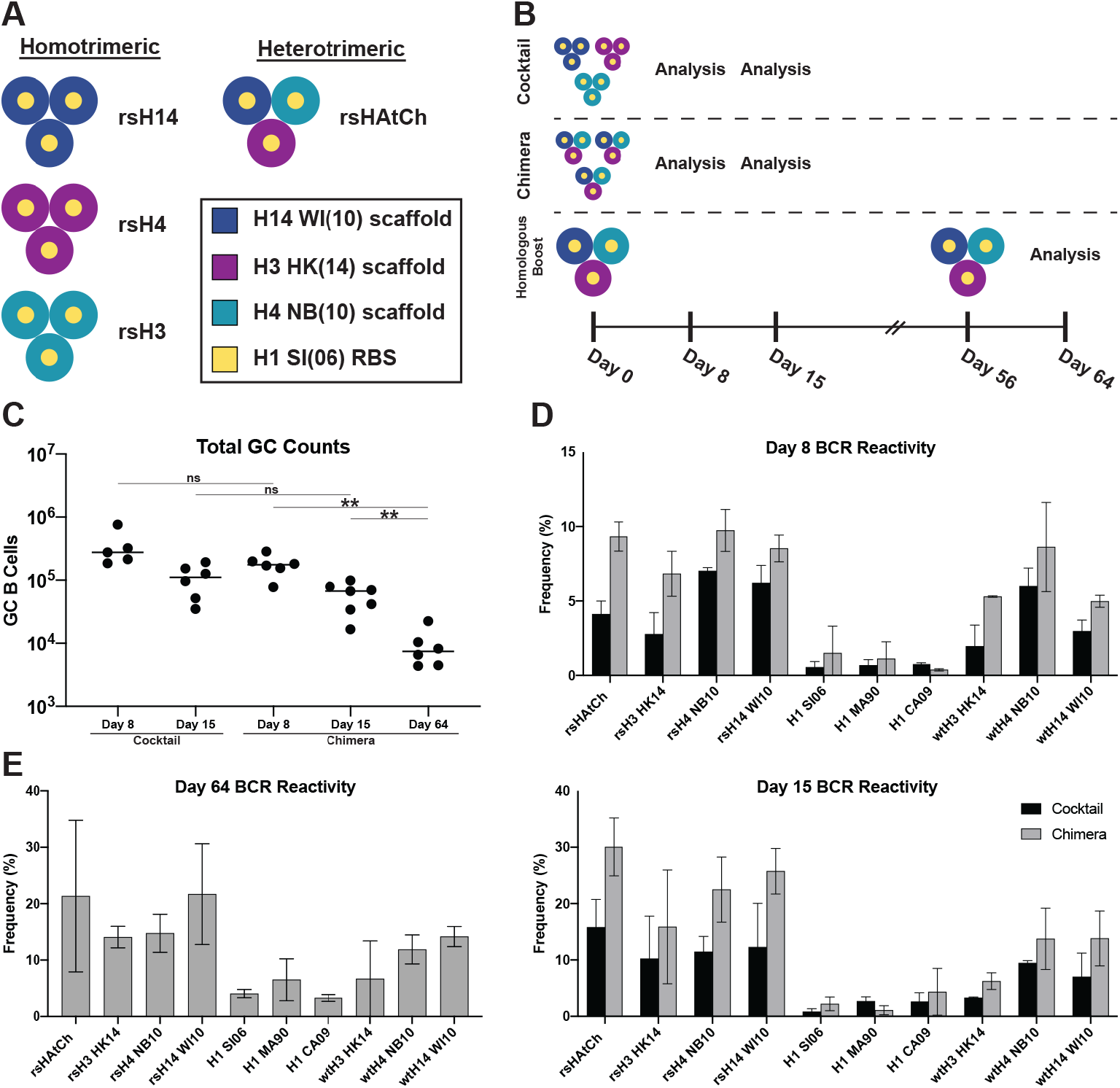
Germinal center analysis of cocktail and chimeric rsHA immunization. **(A)** Cartoon diagrams of homotrimeric rsH14 (blue), rsH4 (purple), and rsH3 (teal), and heterotrimeric rsHAtCh. **(B)** Immunization cohorts and regimens for rsHA homotrimer cocktail, rsHAtCh primary immunization, and rsHAtCh homologous prime-boost immunization. **(C)** Total germinal center B cell populations from each mouse isolated by flow cytometry. **(D)** Combined results of GC analysis for cocktail and rsHAtCh-immunized cohorts at days 8 (top) and 15 (bottom). **(E)** Day 64 GC analysis of rsHAtCh homologous prime-boost immunization cohort.

We determined the overall HA specificity of elicited B cells by screening culture supernatants in a Luminex-based binding assay. The antigen panel included recombinant rsHAtCh, the three rsHAs and their corresponding wild type HAs, as well as three historical H1 HAs. While rsHAtCh recruited a significantly higher frequency of HA^+^ B cells to day 8 GCs than the homotrimer cocktail, the overall frequency was similar between the cohorts by day 15 (**Figure S1A**). Furthermore, the BCR reactivity distributions in day 8 and day 15 GCs were not statistically different between cocktail and rsHAtCh-immunized cohorts. However, we did observe three qualitative trends: 1) response to rsHAs were more prevalent than those to wild type scaffold or H1 HAs, 2) responses to rsH4 and rsH14 components were greater than rsH3, and 3) responses to H1 HA appear to be more prevalent in day 64 GCs than at days 8 or 15 (**Figure 1D-E**). Based on these data, we focused our subsequent analyses on the rsHAtCh cohorts to understand the properties of the elicited B cell repertoires, and specifically, the RBS-directed population.

### rsHAtCh elicits greater frequency of RBS-directed B cells

We next characterized the RBS-directed populations between the cocktail and chimera cohorts to compare binding specificity and potential immune-focusing. To isolate RBS-directed B cells, we designed “ΔRBS” rsHAs with a glycan at Asn190 that abrogates binding to RBS-directed bnAbs (**Figure S1B**) (Lingwood et al., 2012; Whittle et al., 2014). We defined RBS-directed antibodies as those that bound a rsHA, but not the corresponding ΔRBS rsHA or the wild type scaffold HA. While at day 8, there appears to be a relatively higher frequency of RBS-directed B cells in the rsHAtCh cohorts relative to the cocktail, this trend was not statistically significant (**Figure 2A**). However, by day 15, rsHAtCh elicited a significantly greater RBS-directed B cell population. We further classified the RBS-directed responses in terms of “dependency” on the H3, H4, or H14 scaffold periphery surrounding the grafted RBS epitope: we defined “scaffold-dependent” if a B cell bound one rsHA, and “scaffold-independent” if a B cell bound at least two rsHAs. (**Figure 2B**). Based on the H3, H4, and H14 sequence diversity, if a B cell made predominantly peripheral contacts on the grafted RBS epitope, it is more likely to exhibit scaffold dependence. Using these definitions, we find that rsHAtCh elicited a qualitatively higher frequency of RBS-directed B cells that were scaffold independent in both day 8 and day 15 GCs (**Figure 2C-D**). Overall, scaffold dependency in the rsHAtCh cohort shifted from ∼30% scaffold independent at day 8 to ∼60% at day 15, at each point qualitatively outperforming the cocktail immunized cohort (**Figure 2E**). This trend of increasing scaffold independence continued for the rsHAtCh-immunized B cells, reaching ∼70% scaffold independent at day 64. These data suggest that over the course of the GC reaction, rsHAtCh recruits and retains B cells engaging the RBS epitope relative to the cocktail of non-epitope enriched antigens.

**Figure 2:**
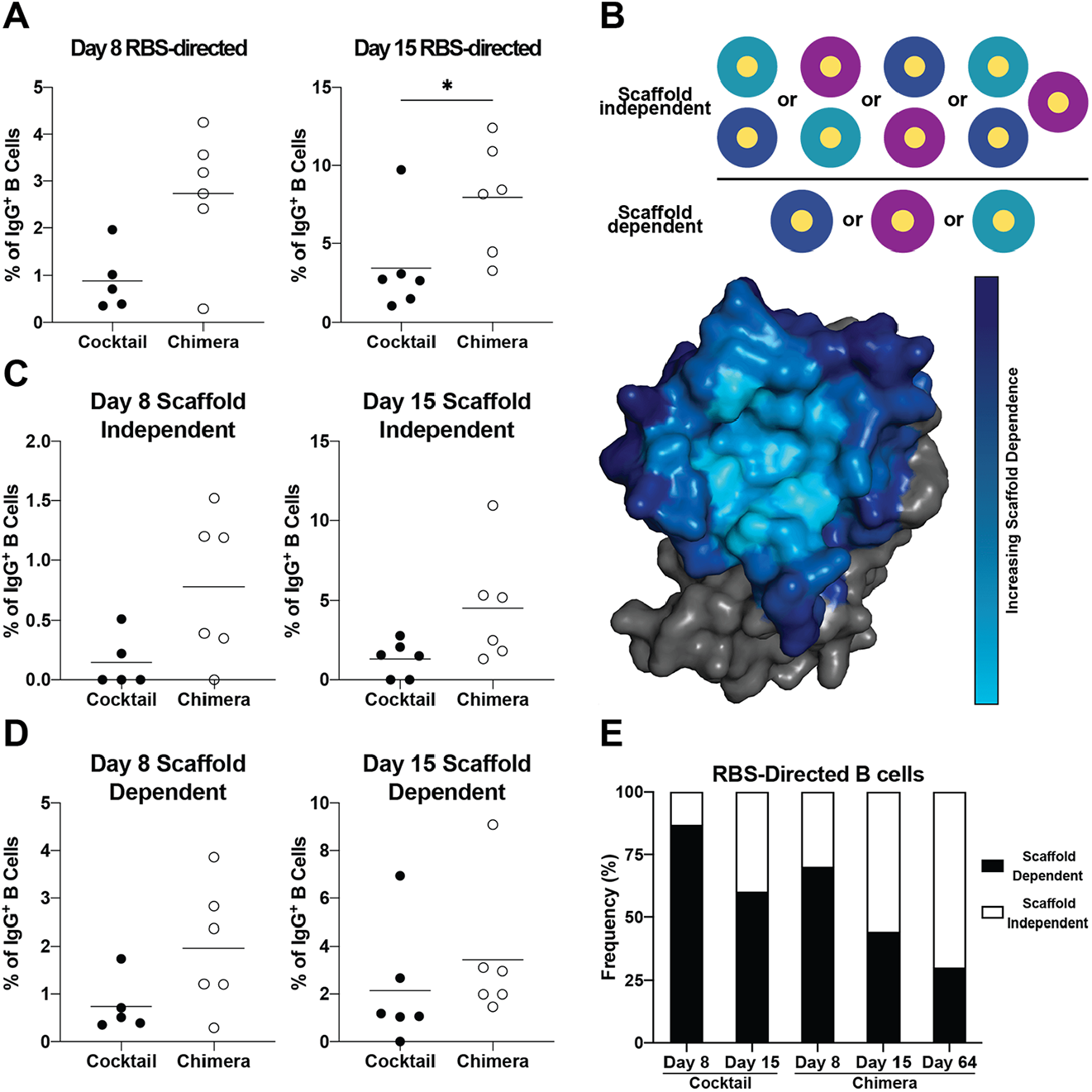
Chimera immunization elicits greater frequency of RBS-directed B cells. **(A)** Frequency of RBS-directed B cells in IgG^+^ Nojima cultures for day 8 (top) and day 15 (bottom) samples. **(B)** Cartoon representation scaffold independent (top) and scaffold dependent (bottom) definitions. Surface representation of the RBS epitope showing increasing likelihood of scaffold dependence with contacts extending away from conserved core residues (**C-D**) Frequency of scaffold independent (top) and scaffold dependent (bottom) antibodies in day 8 and day 15 GCs. **(E)** Balance of scaffold dependent and scaffold independent of RBS-directed B cells pooled across all cohorts. Statistical significance for all comparisons determined using the Mann-Whitney U-test (* denotes p < 0.05).

### Genetic features of rsHAtCh-elicited B cells

We then sequenced rsHAtCh-elicited GC B cells from days 8, 15, and 64 to analyze both the genetic features. Overall, B cells isolated at each time point had a diverse V_H_ gene repertoire (**Figure 3A**). While this diversity is maintained within the RBS-directed B cell population at days 8 and 15, at day 64, we observed an enrichment for B cells with V_H_1-39 or V_H_1-64 genes due to an expansion of clonally related B cells. Notably, >50% of day 64 RBS-directed B cells were V_H_1-39. Day 8 sequences of RBS-directed B cells had a roughly equal distribution of HCDR3 length with an average of 14 amino acids, while days 15 and 64 B cells showed a relative skewing toward 17-18 and 15 amino acid HCDR3s respectively (**Figure 3B**). This observation is consistent with features of canonical human RBS-directed antibodies that have longer HCDR3s necessary to engage the RBS epitope (Ekiert et al., 2012; Krause et al., 2011; Lee et al., 2014; Schmidt et al., 2015b). Of the isolated RBS-directed B cells, the overall levels of V_H_ gene somatic hypermutation (SHM) increased from day 8 to day 15, with a plurality of early and late primary GC B cells containing <1% V_H_ gene mutational frequency; most B cells had <2% mutational frequency overall (**Figure 3C**). In contrast, ∼48% of day 64 GC B cells had >2% SHM, suggesting recall of previously affinity-matured responses. However, 35% of day 64 GC B cells had a <1% mutation frequency, suggesting the presence of *de novo* responses. These data indicate that both the total and RBS-directed populations of rsHAtCh-elicited GC B cells were genetically diverse across day 8, 15, and 64 GCs.

**Figure 3:**
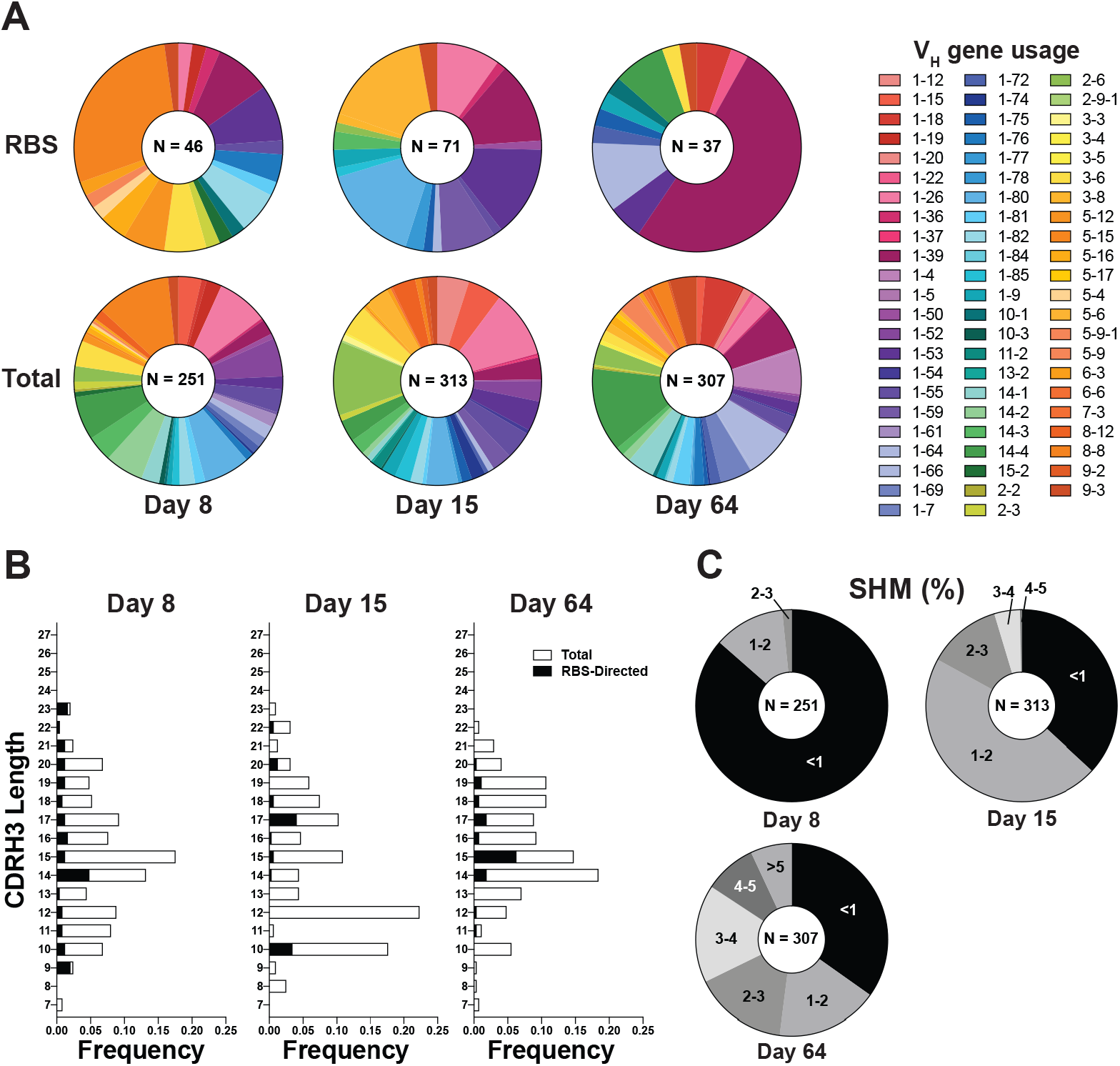
rsHAtCh-elicited germinal centers are diverse. **(A)** V_H_ gene usage of HA^+^ and RBS-directed B cells at day 8 and day 15 timepoints. **(B)** HCDR3 length distributions at day 8, 15, and 64 timepoints for all HA^+^ B cells (white) and RBS-directed B cells (black). **(C)** Somatic hypermutation (SHM) levels in primary (day 8, 15) and secondary (day 64) GCs.

### rsHAtCh elicits clonally related RBS-directed B cells

To further characterize the RBS-directed repertoire, we selected 29 RBS-directed rsHAtCh-elicited antibodies with high avidity in Luminex for one or more HAs in our screening panel for subsequent analysis. 22 clustered into five clonally related populations (CRPs), characterized by identical heavy and light chain gene usages, and similar CDR lengths and sequences (**Figure 4**). We then analyzed heavy and light chain CDRs to identify potential dipeptides (e.g., Val-Asp) that might engage the RBS with sialic acid-like contacts (Schmidt et al., 2015b). 12 RBS-directed B cells from day 15 and 64 GCs had dipeptide motifs in their HCDR3 or LCDR1 loops like dipeptides observed in human RBS-directed broadly neutralizing antibodies (**Figure 4**) (Lee et al., 2014; Schmidt et al., 2015b; Whittle et al., 2011). From day 15 GCs, Ab3, 11, and 23 all contain a Leu/Gly-Asp dipeptide motifs in the HCDR3, while those clonally related to Ab5 or 25, had a reversed Asp-Gly motif. From day 64 GCs, we identified three antibodies with critical dipeptide motifs: Ab42 and 47 have Asp-Gly sequences, while Ab46 has a Glu-Val, positioned in the middle of their 17-19 amino acid HCDR3 loops. Notably, however, not all high affinity RBS-directed antibodies contain sialic acid-like motifs. Antibodies clonally related to htcAb30 (containing V_H_1-39, D_H_2-12, and V_K_4-57 genes) or Ab41 (containing V_H_1-39, D_H_2-4, and V_K_4-57 genes), have long HCDR3 loops from 16 to 18 amino acids without any acidic residues capable of forming sialic acid-like contacts. These data indicate that rsHAtCh immunization can elicit diverse antibodies with similar sequences features of human RBS-directed broadly neutralizing antibodies, and that the elicitation of such RBS-directed populations was observed across multiple mice.

**Figure 4:**
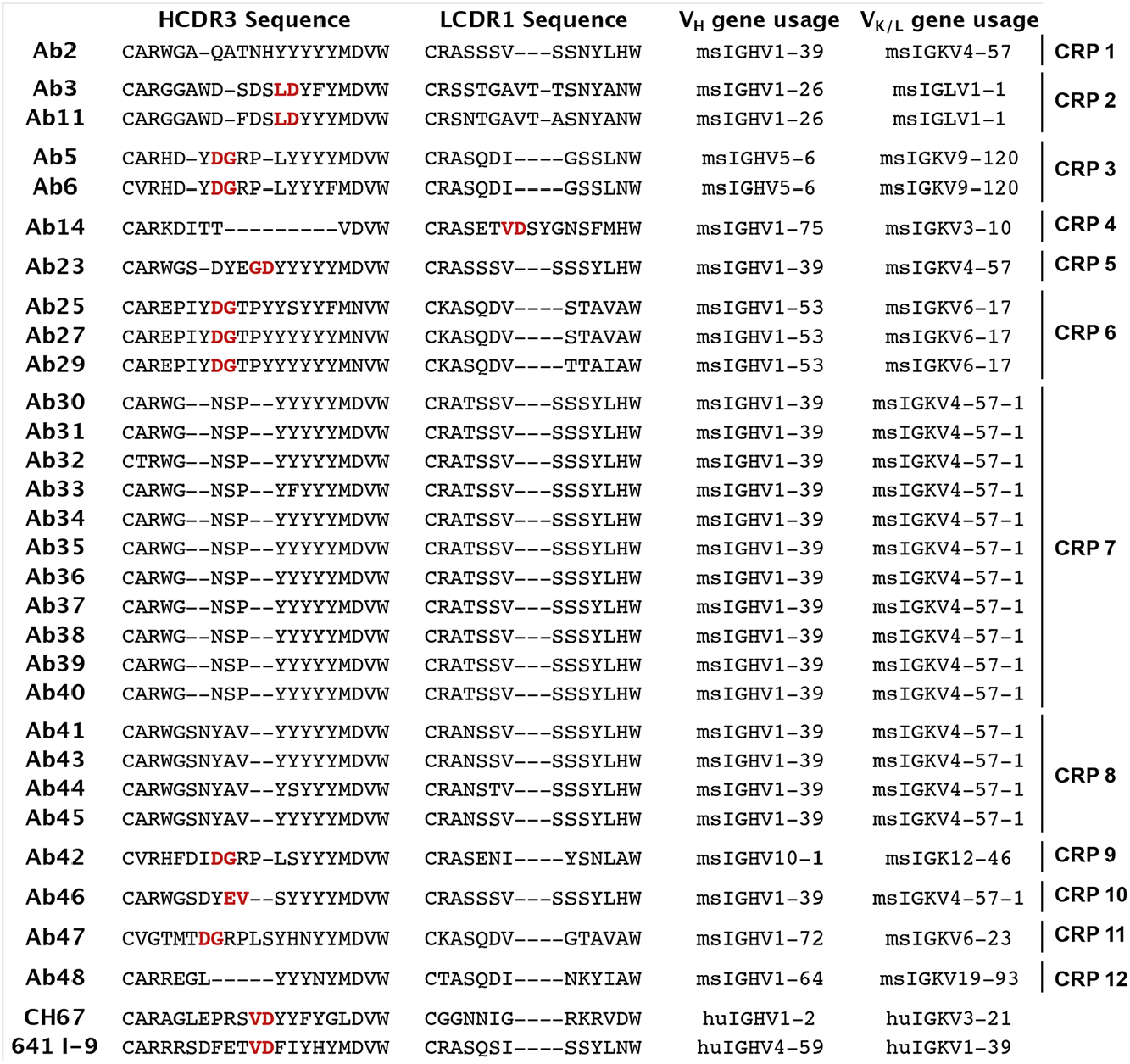
GCs contain clonally-related RBS-directed B cells with critical dipeptide motif. HCDR3 and LCDR1 sequences, and V_H_/V_K/L_ gene usages for high affinity day 15 and day 64 RBS-directed antibodies. Clonally related populations (CRP) from a single mouse are labeled numerically. Critical dipeptide motif for potential sialic acid mimicry shown in red. Dashes denote gaps in the sequence alignment.

### RBS-directed antibodies from secondary GCs bind historical H1s with high affinity

From within the pool of 29 high-avidity RBS-directed antibodies, we subsequently selected the 10 from day 15 GCs, and 11 from day 64 GCs with highest affinity for at least one rsHA to characterize biochemically. We cloned and expressed each as Fabs and tested their affinity against a panel of HAs using biolayer interferometry (BLI). All antibodies were tested against the three rsHA components of rsHAtCh, as well as a panel of historical H1s from 1977 to 2009. 7 of the 10 antibodies from day 15 GCs bound all three rsHAs with <2µM affinity, while the remaining three had detectable affinity for either one or two rsHAs (**Figure 5A**). None of the day 15 GC RBS-directed antibodies had detectable affinity for any historical H1s. To determine if the critical dipeptide motifs we identified by sequence analysis were important for HA binding, we mutated the Asp to Ala in six different antibodies; five of the six dipeptides are in the HCDR3, while for Ab14, the dipeptide is in the LCDR1. These “Δcrit” mutants of Ab3, 11, 25, and 27 abrogated binding to each rsHA (**Figure 5B**). However, Ab14 showed only a ∼10-fold loss of affinity, suggesting that the CDRL1 Asp is involved in antigen binding but not a critical contact. Similarly, Ab23 had a ∼2-fold decrease in affinity for the rsHAs, suggesting that its HCDR3 Asp is not critical for RBS binding.

**Figure 5:**
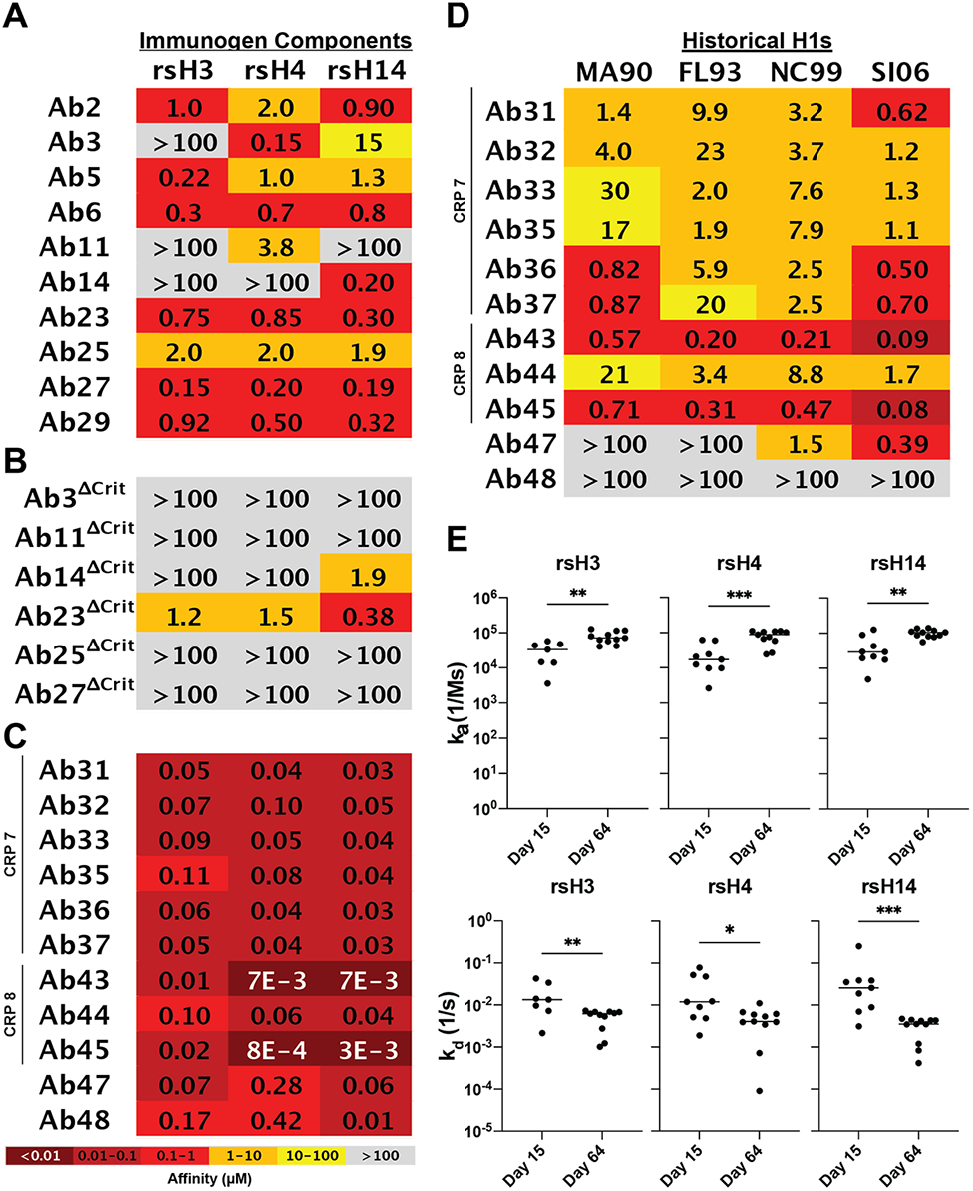
RBS-directed B cells from secondary GCs show increased breadth and affinity. Affinity measurements made using biolayer interferometry (BLI) for **(A)** RBS-directed B cells from day 15 GCs, **(B)** RBS-directed B cells containing Asp→Ala mutations in their critical dipeptide motif, and **(C)** RBS-directed B cells from day 64 GCs against the rsHA components of rsHAtCh. **(D)** Affinity measurements against a panel of historical H1s. **(E)** Association (top) and dissociation (bottom) rates between day 15 and day 64 RBS-directed B cells with highest affinities. Statistical significance was determined using the Mann-Whitney U-test.

In contrast to RBS-directed antibodies from day 15 GCs, those from day 64 GCs were all scaffold independent, engaging the individual rsHA components of rsHAtCh with high affinity (10-100nM) (**Figure 5C**). Furthermore, 10 of the 11 day 64 GC antibodies had high affinity for at least H1 SI-06 and NC-99 (**Figure 5D**). 9 of these antibodies were clonally related to Ab30 or Ab41 and bound historical H1s from 1990 to 2006. Notably, no antibodies bound HA from strains USSR-77, KW-86, BE-95, or pandemic CA-09. Kinetic analysis of day 15 and 64 antibodies showed that the affinity improvement is a consequence of both an increase in the association rate and decrease in the dissociation rate (**Figure 5E**). Thus, breadth against wild type historical H1s can develop after boosting with the rsHAtCh immunogen; these data reflect the overall reactivity profiles observed in the initial Luminex screening (**Figure 1D-E, Figure S1C**).

### Sialic acid-like contacts of rsHAtCh-elicited RBS-directed antibodies

We next obtained high resolution structures of representative rsHAtCh-elicited antibodies that contained putative critical dipeptide motifs or had significant breadth against historical H1s. We determined a co-crystal structure to ∼2.0Å of Ab27 Fab from day 15 GC in complex with the rsH4 head, and a cryoEM structure to ∼3.6Å of Ab36 from day 64 GC in complex with H1 SI-06 (**Figure 6A, D; Figure S5, Table S1-S2**). Structural alignment of wild type H1 SI-06 and rsH4 shows nearly identical secondary structure across the four segments of the grafted RBS epitope (RMSD 0.619Å) (**Figure S4A**). Both antibody footprints engage all four segments of the grafted RBS epitope with predominantly heavy chain contacts and use their HCDR3 and HCDR2 loops to engage the RBS with sialic acid like contacts (**Figure 6B, E**). Ab27 contains an Asp-Gly critical dipeptide motif in the tip of its HCDR3 and makes sialic acid-like contacts with the backbone of Ala137 and Val135 (**Figure 6C**). Ab36 engages the invariant Tyr95 in the RBS core, as well as Gln226 in the 220-loop with its HCDR2 Asn-Tyr motif (**Figure 6F**). Both antibodies make additional HCDR1 contacts with the 220-loop and 190-helix on HA. The HCDR3 loop is positioned above Gly189 and likely explains the lack of binding of Ab31-like or Ab43-like antibodies to H1 BE-95, which has an Arg at this position that would introduce significant steric clash. Both antibodies make multiple germline-encoded contacts with the RBS, which is consistent with the enrichment of V_H_1-53- and V_H_1-39-containing antibodies identified in day 15 and 64 GCs. Notably, the V_H_1-39-containing Ab23 contains the same HCDR2 Asn-Tyr-Gly triad and may make similar RBS contacts, explaining the retained affinity with HCDR3 Δcrit mutations (**Figure S4B**). All biochemically characterized V_H_1-39 RBS-directed Abs contain this Asn-Tyr-Gly motif within the HCDR2.

**Figure 6:**
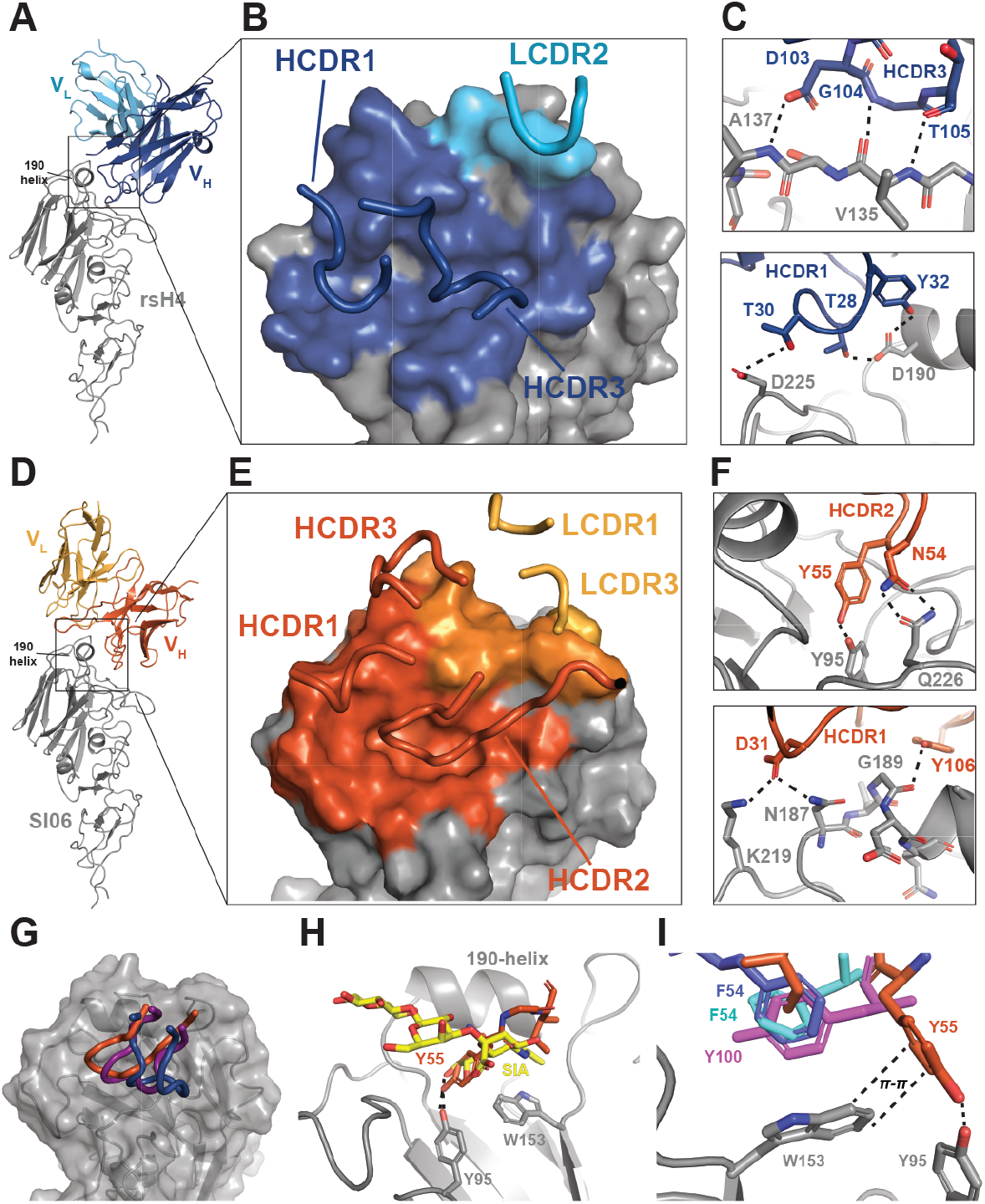
Elicited RBS-directed antibodies exhibit multiple types of receptor mimicry. **(A)** Crystal structure of htcAb27 Fab (top) in complex with rsH4 NB-10 head domain, C_H_ and C_L_ removed for clarity. V_H_ shown in dark blue, V_L_ shown in light blue, and HA shown in grey. **(B)** htcAb27 binding footprint, with contacting CDR loops shown. **(C)** sialic acid-like HCDR3 (top) and HCDR1 (bottom) contacts shown. **(D)** cryoEM structure of htcAb36 in complex with H1 SI06. V_H_ shown in orange, V_L_ shown in yellow, and HA shown in grey. (E) htcAb36 binding footprint, with contacting CDR loops shown. **(F)** sialic acid-mimicking HCDR2 (top) and HCDR1 (bottom) contacts with HA. Salt bridges, hydrogen bonding, and pi-stacking interactions shown as dotted lines. **(G)** Overlay of htcAb27 (blue), htcAb36 (orange), and CH67 (purple) CDRs forming sialic acid-like contacts with HA. **(I)** htcAb36 (orange) and sialic acid (yellow, PDB code 1HGG) contacts with HA. **(J)** π-stacking interaction between HA Trp153 and aromatic residues in htcAb36 (orange), 8M2 (marine blue, PDB code 4HFU), 2G1 (cyan, PDB code 4HG4), and 8F8 (violet, PDB code 4HF5).

Comparison of the RBS-engaging CDRs of Ab27, Ab36, and the canonical sialic acid-mimicking CH67 shows structural convergence despite unique antibody sequences; this is consistent with our previous observations of diverse gene usage in human RBS-directed antibodies (**Figure 6G**) (Schmidt et al., 2015b). Overlay of the HCDR2 from Ab36 with coordinated sialic acid shows striking similarities: the Tyr55 side chain coordinates Tyr95 in HA and occupies roughly the same space as the sialic acid triol moiety, while Ile57 makes van der Waals contacts analogous to the sialic acid acetoamide group (**Figure 6H**). Additionally, Tyr55 π-stacks with the invariant HA Trp153; this interaction has been previously described in a class of human H2-directed receptor-mimicking antibodies containing an Ile-Phe HCDR2 motif or an HCDR3 tyrosine (**Figure 6I**) (Xu et al., 2013). These representative rsHAtCh-elicited antibodies from day 15 and 64 GCs show that enriched clonally related RBS-directed antibodies have similar structural features as human RBS-directed bnAbs.

## DISCUSSION

Here we show that increasing the frequency of the RBS epitope on a single immunogen led to greater expansion of RBS-directed B cells in primary GCs, with RBS-directed subpopulations in both primary and secondary GCs containing sialic acid-like CDR moieties. Indeed, structures of representative RBS-directed antibodies showed two distinct mechanisms of receptor mimicry, both of which have been previously characterized in human broadly neutralizing RBS-directed antibodies. These data indicate that immunization with an RBS-enriched focuses to the RBS while maintaining a high degree of genetic and structural diversity comparable to what has been seen in human RBS-directed responses (Lee et al., 2014; Lee et al., 2012; Schmidt et al., 2015b; Xu et al., 2013).

Recognition of a broadly conserved epitope by an antibody does not necessarily confer breadth (Zost et al., 2019). Indeed, the breadth of RBS-directed antibody unmutated common ancestors (UCAs) isolated from human donors is often quite limited; breadth is acquired through multiple rounds of antigen exposure and subsequent affinity maturation (Schmidt et al., 2015a). Repeated exposure to rsHAtCh led to increased RBS-directed antibody breadth, with over 40% of RBS-directed Abs engaging H1s across an almost two-decade range. Lack of reactivity of Ab30- or Ab41-like antibodies to H1 BE-95, is likely due to a steric clash with Arg189; the other historical strains have a Gly at this position. Notably, rsHAtCh-elicited antibodies did not engage the pandemic H1 CA-09; this is likely due to the significant sequence difference between the grafted H1 RBS epitope based on the 2006 strain (Bajic et al., 2020). Sequential immunization with a series of RBS-enriched immunogens presenting antigenically drifted RBS grafts might maintain core RBS enrichment while allowing for the gradual broadening of antibody reactivities to develop both retrospective and prospective breadth across distinct antigenic clusters.

How epitope enrichment may influence patterns of immunodominance is presumably multifactorial, but is likely driven by the enhanced avidity between BCRs and the grafted H1 RBS epitope on the rsHAtCh immunogen; avidity is critical during cell recruitment into GCs, antigen acquisition in the GC light zone, and reactivation of B cells from memory (Finney et al., 2018; Viant et al., 2020; Yeh et al., 2018). A computational model based on the data presented here, including the qualitative observation that rsHACh more readily recruit B cells to GCs, provides a potential mechanistic explanation for our observations: namely that the rsHAtch binds cross-reactive BCRs multivalently and therefore can capture more antigen early in the GC and that when T cell selection is a stringent constraint, it favors the evolution of broad reactivity (Yang et al., 2022). This computational study shows how rational immunogen design can result in a synergistic effect between antigen capture by B cells and the stringency of T cell selection in the GC for developing cross-reactive B cells. The data reported here, and in the accompanying computational study, further our understanding of fundamental immunological questions. The extent to which secondary GCs are reseeded by memory B cells following identical antigen exposures is not entirely clear (Pape and Jenkins, 2018; Shlomchik, 2018); this is relevant to our homologous rsHAtCh immunization, here. While we observed accumulation of additional somatic mutations in day 64 GC B cells, a plurality of B cells contained a V_H_ mutation frequency of <1%, suggesting both the recruitment and maturation of antigen-experienced B cells as well as *de novo* recruitment of naïve B cells. However, the development of H1 breadth among multiple clonally-related populations from the rsHAtCh immunizations of RBS-directed antibodies supports a model in which a subset of B cells are recalled and further affinity matured upon re-exposure.

These experiments are complementary to our rsHAtCh studies in IGHV1-2 HC2 KI mice (Caradonna et al., 2022); where antibodies use the IGHV1-2 sequence but have human-like diversity in the HCDR3 loop, which is the principal source of BCR diversity (Sangesland et al., 2019; Sangesland et al., 2020). In that study, we analyzed the serum and class-switched B cell compartments of heterologous-boosted mice primed with H1 SI-06 as a surrogate for immune imprinting. While we observed a significant expansion of H1 RBS-directed B cells after rsHAtCh immunization across both compartments, the repertoire was both limited in breadth and genetically restricted. This contrasts with our observations here, where homologous-boosted J_H_6 mice, in the absence of immune imprinting to a specific HA, resulted in significant breadth to historical H1 HAs. Importantly, these two studies examined different immune compartments, primary and secondary GCs versus serum and memory compartments, and thus the timing of analyses are necessarily different. Understanding how immune imprinting influences subsequent recall of responses in the GC and serum compartments, ultimately targeting conserved sites elicited by rationally designed immunogens is necessary.

Current immunogen design strategies aim to focus antibody responses through removing or occluding ‘off-target’ competing responses, such as stem-only nanoparticle or hyperglycosylated HA immunogens (Bajic et al., 2019; Corbett et al., 2019; Yassine et al., 2015). The removal or shielding of immunodominant HA head epitopes reduces the extent of inter-clonal competition but does not confer any advantage to a specific subset of B cells. In contrast, heterologous antigen display is thought to confer an avidity advantage to GC B cells recognizing cross-reactive epitopes due to the asymmetric conservation of epitopes when distinct antigens are present (Boyoglu-Barnum et al., 2021; Kanekiyo et al., 2019). However, not all mosaic nanoparticle designs are superior to cocktail immunizations at eliciting broadly neutralizing antibodies, suggesting that certain patterns of epitope enrichment may be more favorable than others, and that heterologous display can be further optimized for specific epitopes (Cohen et al., 2021). Our engineered rsHAtCh immunogen, enriched for the H1 RBS epitope, offers a putative competitive advantage directed towards RBS-directed B cells. The trajectory of RBS-directed antibodies progressively shifting towards scaffold independence across the GC reaction further supports this notion, as does the expansion of RBS-directed B cells with affinity for historical H1s.

While prior immunogens in the field were designed to elicit immune responses through heterologous HA display, this was in the context of validating potential vaccine candidates. As such, cross-reactivity, neutralization, and protection, independent of epitope, were the primary objectives. Here, our rsHAtCh immunogen explicitly tests the underlying principle suggested by these preclinical validation studies; it analyzes the biology of heterologous display optimized for a single subdominant epitope, and shows that epitope-enriched immunogens can enhance specific responses against single epitopes. Thus, epitope enrichment can be used with other immunogen design strategies, and more broadly adapted to other conserved and protective epitopes.

## LIMITATIONS OF THE STUDY

A primary limitation of this study was sample size, given the significant stochasticity of the germinal center our observations were more qualitative and we do not intend to overstate our findings. Binding measurements with Luminex-based assays or BLI may not fully represent *in vivo* avidities due monomeric versus multimeric display of BCRs on the cell surface. Conceptually, the ways in which increased an epitope-enriched immunogen may impact the trajectory of the GC reaction is highly complex, including among other components, B cell recruitment to germinal centers, retention in germinal centers, antigen acquisition, stronger T cell help, and reactivation of anamnestic responses. The deconvolution of these processes will require further fundamental study.

## STAR METHODS

### RESOURCE AVAILABILITY

#### Lead Contact

Further information and requests for resources and reagents should be directed to and will be fulfilled by the lead contact, Aaron G. Schmidt.

#### Materials Availability

All unique/stable reagents generated in this study will be made available on request, but we may require a payment and/or a completed materials transfer agreement if there is potential for commercial application. For non-commercial use, all unique/stable reagents generated in this study are available from the lead contact with a completed materials transfer agreement.

#### Data and Code Availability

- CryoEM data are deposited in the Electron Microscopy Data Base and are available as of the date of publication. X-ray crystallography are deposited in the Protein Data Bank (7TRZ) and are available as of the date of publication. B cell receptor sequences are deposited in Genbank: accession numbers are listed in the key resources table.
- This paper does not report original code.
- Any additional information required to reanalyze the data reported in this paper is available from the lead contact upon request.

### EXPERIMENTAL MODEL AND SUBJECT DETAILS

#### Cell Lines

FreeStyle 293F cells (Thermo Fisher Cat#R79007; RRID: CVCL_D603, female) and Expi293F cells (Thermo Fisher Cat#A14527; RRID: CVCL_D615, female) were cultured following manufacturer’s instructions. High Five cells (BTI-TN-5B1-4, female) (*Trichoplusia ni*) and Sf9 cells (*Spodoptera frugiperda*, female) were cultured in EX-CELL 405 SFM (Sigma, Cat# 14405C) and Sf-900 II SFM (Gibco, Cat# 10902096) respectively in accordance with the manufacturer’s instructions.

#### Mouse Model

The huJ_H_6 KI mice (Bajic et al., 2020) were bred and maintained under specific pathogen-free conditions at the Duke University Animal Care Facility. Eight to 12-week-old mice were used for immunizations. Mouse immunization experiments were approved by the Duke University Institutional Animal Care and Use Committee.

## METHOD DETAILS

### Immunogen and Luminex Protein Expression and Purification

rsH3, rsH4, and rsH14 HA head constructs were derived from A/long-tailed duck/Wisconsin/ 10OS3918/2010 (H14 WI-10), A/American black duck/New Brunswick/00464/2010(H4 NB-10), and A/HongKong/4801/2014 (H3 HK14) scaffolds resurfaced with the H1 SI-06 RBS described previously (Bajic et al., 2020). All constructs were codon optimized by Integrated DNA Technologies and purchased as gblocks that were then cloned into pVRC and sequence confirmed via Genewiz. To generate the rsHAtCh construct, each rsHA was subcloned into a pVRC protein expression vector containing individual components of the non-collagenous domain 2 (NC2)-derived hetero-trimerization motif. Each component containing a rsHA, hetero-trimerization tag, and either 8xHis, FLAG, or streptavidin binding protein (SBP) tag, was amplified via PCR to introduce overlapping P2A sequence regions, and all three fragments were combined with linearized pVRC vector via Gibson assembly into the final expression construct. Trimeric constructs for Luminex screening contained C-terminal HRV 3C-cleavable 8xHis and SBP tags.

Expi 293F cells (ThermoFisher) or HEK293F cells (ThermoFisher) were used to express proteins. Transfections were performed with Expifectamine or PEI reagents per the manufacturer’s protocol. For homotrimeric HA and Luminex antigens, transfections were harvested and centrifuged for clarification after 5-7 days. Cobalt-TALON resin (Takara) was used to perform immobilized metal affinity chromatography via the 8xHis tag. Proteins were eluted using imidazole, concentrated, and passed over a Superdex 200 Increase 10/300 GL (GE Healthcare) size exclusion column. Size exclusion chromatography was performed in 10mM Tris pH 7.5, 150mM NaCl. For immunogens, HRV 3C protease (ThermoScientific) cleavage of affinity tags was performed prior to immunization. Cobalt-TALON resin was used for a repurification to remove the His-tagged HRV 3C protease, cleaved tag, and remaining uncleaved protein.

rsHAtCh protein was expressed via transient transfection in HEK293F cells for 5 days, and all subsequent purification steps were carried out at 4°C. Clarified culture supernatants were purified over Cobalt-TALON affinity resin (Takara), with subsequent purification via size exclusion chromatography (SEC) over a Superdex 200 Increase (Cytiva) column equilibrated in 10mM Tris, 150mM NaCl, pH 7.5 running buffer. Fractions containing only trimeric rsHAtCh were then passed over G1 anti-FLAG-agarose resin (Genscript) equilibrated in 10mM Tris, 150mM NaCl, pH 7.5, incubated for 1 hour rotating, and eluted with 50mM Tris, 150mM NaCl, containing 160µg/mL 1xFLAG peptide (ApexBio) pH 7.5 following a 1-hour incubation with mild agitation. The sample was then purified a third time over Streptavidin-agarose resin (Pierce) preequilibrated in 10mM Tris, 150mM NaCl, pH 7.5, incubated for 1 hour rotating, washed with 20mM HEPES, 150mM NaCl, 5% glycerol, pH 7.0, and eluted with 20mM HEPES, 150mM NaCl, 4mM biotin, 5% glycerol pH 7.0 after a 1-hour incubation. All affinity and heterotrimerization tags were removed using HRV 3C protease (Thermo), and the sample was purified a final time via SEC equilibrated in PBS, and the trimeric peak was harvested and concentrated to 2.0mg/mL for immunizations.

### Fab and IgG Expression and Purification

The variable heavy and light chain genes for each antibody were purchased as codon optimized gBlocks by Integrated DNA Technologies, and subsequently cloned into pVRC constructs containing the appropriate constant domains as previously reported (Schmidt et al., 2015a; Schmidt et al., 2015b). The Fab heavy chain vector contained a HRV 3C-cleavable 8xHis tag, and the IgG heavy chain vector contained HRV 3C-cleavable 8xHis and SBP tags. The same transfection and purification protocol as used for the immunogens and coating proteins was used for the Fabs and IgGs.

### Biolayer Interferometry

Biolayer interferometry (BLI) experiments were performed using a BLItz instrument (Fortebio) with Ni-NTA biosensors (Fortebio). All proteins were diluted in 10mM Tris, 150mM NaCl, pH 7.5. HA proteins were immobilized to the biosensors, and Fabs were used as the analytes. To determine binding affinities, single-hit measurements were performed starting at 10 *μ*M to calculate an approximate *K*_*D*_ in order to evaluate which concentrations should be used for subsequent titrations. Measurements at a minimum of four additional concentrations were performed. Vendor-supplied software was used to generate a final *K*_*D*_ estimate via a global fit model with a 1:1 binding isotherm. All binding studies were carried out in 10mM Tris, 150mM NaCl, pH 7.5 at room temperature.

### Mouse Strains and Murine Immunizations

The huJ_H_6 KI mice (Bajic et al., 2020) were bred and maintained under specific pathogen-free conditions at the Duke University Animal Care Facility. Eight to 12-week-old mice were immunized with 20 *μ*g of rsHAtCh antigens or an equimolar amount of an rsHA cocktail (rsH3, rsH4, and rsH14) in the footpad of the right hind leg. Eight weeks later, cohort of mice were boosted with 20 *μ*g rsHAtCh in the hock ipsilaterally. All antigens were mixed with Alhydrogel^®^ before immunizations. Mouse immunization experiments were approved by the Duke University Institutional Animal Care and Use Committee.

### Flow Cytometry

Single cell suspensions were obtained from LN cells and labeled with mAbs to identify GC B cells (GL-7+/B220*hi*/CD38*lo*/IgD-/CD138-) as described (Kuraoka et al., 2016). Labeled cells were analyzed/sorted in a FACS Vantage with DIVA option (BD Bioscience). Flow cytometric data were analyzed with FlowJo software (Treestar Inc.). Doublets were excluded by FSC-A/FSC-H gating strategy. Cells that take up propidium iodide were excluded from our analyses.

### Nojima Culture

Single B cells were cultured in the presence of NB-21.2D9 feeder cells (Kuraoka et al., 2016). One day before B-cell sorting, NB-21.2D9 cells were seeded into each well of 96-well plates (2,000 cells/well) in B cell media (BCM); RPMI-1640 (Invitrogen) supplemented with 10% HyClone FBS (Thermo scientific), 5.5 × 10^−5^ M 2-mercaptoethanol, 10 mM HEPES, 1 mM sodium pyruvate, 100 units/ml penicillin, 100 *μ*g/ml streptomycin, and MEM nonessential amino acid (all Invitrogen). Next day (Day 0), recombinant mouse IL-4 (Peprotech; 2 ng/ml) was added to the cultures, and then single B cells were directly sorted into each well using a FACS Vantage. On day 2, 50% (vol.) of culture media were removed from cultures and 100% (vol.) of fresh BCM were added to the cultures. On days 3 to 8, two-thirds of the culture media were replaced with fresh BCM every day. On day 10, culture supernatants were harvested for ELISA determinations and culture plates that contain cell pellets of expanded, clonal B cells were stored at -80°C for V(D)J amplifications.

### ELISA and Luminex Multiplex Assay

IgG^+^ culture supernatants determined by standard ELISA were screened for the reactivity to rHAs by Luminex multiplex assay (Kuraoka et al., 2016; Watanabe et al., 2019). Briefly, Culture supernatants were diluted (1: 10) in 1×PBS containing 1% BSA, 0.05% NaN3 and 0.05% Tween20 (assay buffer) with 1% milk and incubated for 2 hours with the mixture of antigen-coupled microsphere beads in 96-well filter bottom plates (Millipore). After washing with assay buffer, beads were incubated for 1 hour with PE-conjugated goat anti-mouse IgG Abs (Southern Biotech). After three washes, the beads were re-suspended in assay buffer and the plates were read on a Bio-Plex 3D Suspension Array System (Bio-Rad). The following antigens were coupled with carboxylated beads (Luminex Corp): BSA (Affymetrix), goat anti-mouse Igκ, goat anti-mouse Igλ, goat anti-mouse IgG (all Southern Biotech), and a panel of rHAs as listed in the rHA expression methods subsection.

### BCR Sequencing

Rearranged V(D)J gene sequences were obtained from single-cell cultured GC B cells by a nested PCR as described (Kuraoka et al., 2016; Rohatgi et al., 2008; Tiller et al., 2009) with modifications. Total RNA was extracted from selected samples using Quick-RNA 96 kit (Zymo Research). cDNA was synthesized using Superscript III with 0.2 *μ*M each of gene-specific reverse primers (mIgGHGC-RT, mIgKC-RT, mIgLC23-RT, mIgLC1-RT, and mIgLC4-RT) at 50°C for 50 min followed by 85°C for 5 min. cDNA samples were then subjected to two rounds of PCR using Herculase II fusion DNA polymerase (Agilent Technologies).

In some experiments, cDNA was synthesized from DNase I-treated RNA using SMARTScribe™ Reverse Transcriptase (Clontech) with 0.2 *μ*M each of gene-specific reverse primers (same as above) and 1 *μ*M of 5’ SMART template-switching oligo (TSO-bioG). cDNA was then subjected to two rounds of SMART PCR using Herculase II fusion DNA polymerase with a common forward primer (5Anchor1-FW1) for both the 1^st^ and the 2^nd^ PCR, and reverse primers.

V(D)J amplicands were submitted to Duke DNA sequencing facility to obtain DNA sequences. The rearranged V, D, and J gene segments were first identified using IMGT/V-QUEST (http://www.imgt.org/) or Cloanalyst (Kepler, 2013), and then numbers and kinds of point mutations were determined.

IMGT High V-Quest was used to analyze variable heavy and light chain sequences.

### CryoEM Grid Preparation and Image Recording

Complexes of H1N1 A/Solomon Island/3/2006 Q226 hemagglutinin (HA) with ab36 were formed by combining at a final concentration 2.5 mg/mL HA with 1.8 mg/mL Fab (2-fold excess of binding sites) in a buffer composed of 10 mM Tris pH 7.5 with 150 mM NaCl. Octyl β-D-glucoside (OG) was added at a final concentration of 0.08% (w/v) to correct the HA orientation bias. HA°Fab complexes were incubated for 30 minutes on ice before application to C-flat 1.2-1.3 400 Cu mesh grids (Protochips). Grids were glow discharged (PELCO easiGlow) for 30 seconds at 15 mA and prepared with a Gatan Cryoplunge 3 by applying 3.5 uL of sample and blotting for 4.0 seconds in the chamber maintained at a humidity between 86% and 90%.

### CryoEM Image Analysis and 3D Reconstruction and Model Fitting

Images for HA and Ab36 Fab complexes were recorded on a Titan Krios microscope operated at 300 keV with a Gatan BioQuantum GIF/K3 direct electron detector. Automated image acquisition was performed with Serial EM. Specifications and statistics for images are reported in **Table S2**.

Image analysis was carried out in RELION. Beam-induced motion correction of micrograph movies was performed with UCSF MotionCor2 followed by contrast transfer function estimation with CTFFIND-4.1, both as implemented in RELION. Particles were picked from motion corrected micrographs by a general model using crYOLO. Extracted particles were down sampled by a factor of 3 and subjected to 2D classification. Views were primarily down and near the 3-fold axis. An initial C3-symetric 3D model was prepared from small subset of particle that suffered from a preferred orientation. A subset of 2D classes exhibited side views, however, they consisted of a dimer of HA and Fab complexes. These particles were separated from the rest and re-extracted with symmetric offsets to create two HA-centered particle stacks. These side views were combined with the top views and subjected to 3D classification and refinement yielding a Nyquist-limited reconstruction. Re-extraction of particles at the full pixel size and subsequent 3D classification and refinement were performed with a mask that omitted the stem of HA and the Fc of the Fab to yield a 4.2 Å map. CTF refinement was performed to correct beam tilt, trefoil, anisotropic magnification, and per particle defocus in RELION. Bayesian polishing was also performed in RELION leading to a 3.7 Å reconstruction following 3D refinement. The final map was improved to 3.6 Å resolution with B-factor sharpening (**Figure S2**).

Heavy and light chains of PDB entries 4KUZ and 1FIG were aligned and extracted to make an initial model for the Fab. A protomer of H1 Solomon Islands was extracted from PBD entry 5UGY. These initial models were ridged body fit into the Ab36 cryoEM density map using Chimera. The gross positions of the initial models were further refined by ridged body real-space refinement with Phenix. The sequences of the initial antibody Fv domains were corrected to match the actual sequence in COOT. The coordinates were further real space refined with Phenix. Iterative cycles of model adjustment and refinement were performed to eliminate Ramachandran outliers and non-covalent interatomic distances of less than 2.2A. Map-to-model statistics and FSC curves were prepared with Phenix (**Table S2** and **Figure S2**).

### Crystallization

rsH4 (H4 A/American black duck/New Brunswick/00464/2010 with H1 A/Solomon Islands/03/ 2006 grafted RBS) HA1 head domain and htcAb27 Fab were incubated at a 1:1.3 molar ratio and complexes were purified using a Superdex 200 gel filtration column equilibrated in 10 mM Tris-HCl, 150 mM NaCl, pH 7.5. Complexes were concentrated to 16 mg/mL and crystals were grown at 18° via hanging drop vapor diffusion. htcAb27 and rsH4 head complex crystals were grown in 12% PEG 20,000, 100 mM PIPES, 100 mM NaCl, pH 6.0. Crystals were cryoprotected in mother liquor supplemented with 10% 2-methyl-2,4-pentanediol (MPD) and flash frozen in liquid nitrogen.

### Structure Determination and Refinement

Diffraction data were recorded at 0.999 Å with a rotation of 1° per image on the 24-ID-C beamline at the Advanced Photon Source, Argonne National Laboratory. Data was indexed and reduced by the RAPD data-processing pipeline (https://github.com/RAPD/RAPD), using XDS (Kabsch, 2010) for indexing and integration and the CCP4 programs AIMLESS and TRUNCATE (Winn et al., 2011) for scaling and structure-factor calculation. The data were scaled in the P2_1_2_1_2_1_ space group, and an initial structure was determined using molecular replacement (MR) in PHASER using the first generation rsH4 NB-10 HA head (PDB 6UR5) for the HA head search model, and separate models for the V_H_V_L_ (PDB 1A6U, 5I66 respectively) and C_H_C_L_ (PDB 1FIG). The additional mutations in the grafted epitope of the rsH4 NB-10 v2 HA head were rebuilt from the starting model. The structure was then refined with phenix.refine using reciprocal-space, real-space, grouped B factor, and Translation Libration Screw-rotation (TLS) refinement (McCoy et al., 2007). There was limited density for several residues on the N and C termini of the HA construct, which did not improve during refinement and were removed from the final model. All model building was performed in Coot (Emsley and Cowtan, 2004), and model quality was assessed using the MolProbity server (Chen et al., 2010). Figures were created using PyMOL Molecular Graphics System (v2.4.1, Schrödinger LLC). Refinement statistics are shown in Table S1. X-ray crystallography data were deposited in the Protein Data Bank (7TRZ)

## QUANTIFICATION AND STATISTICAL ANALYSIS

Curve fitting and statistical analyses were performed with GraphPad Prism (version 9). Non-parametric statistics were used throughout where feasible. Tests, numbers of animals, and statistical comparison groups are indicated in each of the Figure Legends. The Mann-Whitney U test was used to compare two populations without consideration for paired samples. A p value < 0.05 was considered significant.

## ACKNOWLEDGMENTS

We thank Stephen Harrison, Jared Feldman, Kevin R. McCarthy, and Goran Bajic for helpful discussion. We thank Xiaoe Liang at Duke University for assistance and mouse colony maintenance. This work is based upon research conducted at the Northeastern Collaborative Access Team beamlines, which are funded by the National Institute of General Medical Sciences from the National Institutes of Health (P30 GM124165). This research used resources of the Advanced Photon Source, a U.S. Department of Energy (DOE) Office of Science User Facility operated for the DOE Office of Science by Argonne National Laboratory under Contract No. DE-AC02-06CH11357. We thank the beamline staff at NE-CAT for help. We acknowledge support from the NIH for R01AI146779 (A.G.S.), P01AI89618-A1 (A.G.S, M.K.) and T32 GM007753 (T.M.C.). This research has been funded in whole or part with federal funds under a contract from the National Institute of Allergy and Infectious Diseases, NIH contract 75N93019C00050.

## AUTHOR INFORMATION

### Author Contributions

T.M.C., A.W., G.K., M.K., A.G.S designed research; T.M.C., I.W.W., A.A.R., S.S., A.W., M.K., A.G.S., performed research; T.M.C., I.W.W., A.A.R., S.S., A.W., G.K., M.K., A.G.S. analyzed data; T.M.C and A.G.S wrote the paper; T.M.C., I.W.W., A.A.R., S.S., A.W., G.K., M.K., A.G.S. edited and commented on the paper.

### Correspondence and requests for materials should be addressed to

Aaron G. Schmidt or Masayuki Kuraoka

## REFERENCES

Andrews, S.F., Huang, Y., Kaur, K., Popova, L.I., Ho, I.Y., Pauli, N.T., Dunand, C.J.H., Taylor, W.M., Lim, S., Huang, M., et al. (2015). Immune history profoundly affects broadly protective B cell responses to influenza. Science Translational Medicine 7, 316ra192–316ra192.

Angeletti, D., Gibbs, J.S., Angel, M., Kosik, I., Hickman, H.D., Frank, G.M., Das, S.R., Wheatley, A.K., Prabhakaran, M., Leggat, D.J., et al. (2017). Defining B cell immunodominance to viruses. Nature immunology 18, 456–463.

Bajic, G., Maron, M.J., Adachi, Y., Onodera, T., McCarthy, K.R., McGee, C.E., Sempowski, G.D., Takahashi, Y., Kelsoe, G., Kuraoka, M., et al. (2019). Influenza Antigen Engineering Focuses Immune Responses to a Subdominant but Broadly Protective Viral Epitope. Cell Host & Microbe 25, 827-835.e826.

Bajic, G., Maron, M.J., Caradonna, T.M., Tian, M., Mermelstein, A., Fera, D., Kelsoe, G., Kuraoka, M., and Schmidt, A.G. (2020). Structure-Guided Molecular Grafting of a Complex Broadly Neutralizing Viral Epitope. ACS Infectious Diseases 6, 1182–1191.

Boyoglu-Barnum, S., Ellis, D., Gillespie, R.A., Hutchinson, G.B., Park, Y.-J., Moin, S.M., Acton, O.J., Ravichandran, R., Murphy, M., Pettie, D., et al. (2021). Quadrivalent influenza nanoparticle vaccines induce broad protection. Nature.

Caradonna, T.M., Ronsard, L., Yousif, A., Windsor, I.W., Hecht, R., Bracamonte-Moreno, T., Roffler, A.A., Maron, M.J., Feldman, J., Marchiori, E., et al. (2022). An epitope-enriched immunogen expands responses to a conserved viral site. Submitted.

Chen, V.B., Arendall, W.B., III, Headd, J.J., Keedy, D.A., Immormino, R.M., Kapral, G.J., Murray, L.W., Richardson, J.S., and Richardson, D.C. (2010). MolProbity: all-atom structure validation for macromolecular crystallography. Acta Crystallographica Section D 66, 12–21.

Cohen, A.A., Yang, Z., Gnanapragasam, P.N.P., Ou, S., Dam, K.-M.A., Wang, H., and Bjorkman, P.J. (2021). Construction, characterization, and immunization of nanoparticles that display a diverse array of influenza HA trimers. PloS one 16, e0247963.

Corbett, K.S., Moin, S.M., Yassine, H.M., Cagigi, A., Kanekiyo, M., Boyoglu-Barnum, S., Myers, S.I., Tsybovsky, Y., Wheatley, A.K., Schramm, C.A., et al. (2019). Design of Nanoparticulate Group 2 Influenza Virus Hemagglutinin Stem Antigens That Activate Unmutated Ancestor B Cell Receptors of Broadly Neutralizing Antibody Lineages. mBio 10.

Corti, D., Voss, J., Gamblin, S.J., Codoni, G., Macagno, A., Jarrossay, D., Vachieri, S.G., Pinna, D., Minola, A., Vanzetta, F., et al. (2011). A neutralizing antibody selected from plasma cells that binds to group 1 and group 2 influenza A hemagglutinins. Science (New York, NY) 333, 850–856.

Ekiert, D.C., Kashyap, A.K., Steel, J., Rubrum, A., Bhabha, G., Khayat, R., Lee, J.H., Dillon, M.A., O’Neil, R.E., Faynboym, A.M., et al. (2012). Cross-neutralization of influenza A viruses mediated by a single antibody loop. Nature 489, 526–532.

Emsley, P., and Cowtan, K. (2004). Coot: model-building tools for molecular graphics. Acta Crystallographica Section D 60, 2126–2132.

Finney, J., Yeh, C.-H., Kelsoe, G., and Kuraoka, M. (2018). Germinal center responses to complex antigens. Immunol Rev 284, 42–50.

Hai, R., Krammer, F., Tan, G.S., Pica, N., Eggink, D., Maamary, J., Margine, I., Albrecht, R.A., and Palese, P. (2012). Influenza Viruses Expressing Chimeric Hemagglutinins: Globular Head and Stalk Domains Derived from Different Subtypes. Journal of virology 86, 5774.

Harshbarger, W.D., Deming, D., Lockbaum, G.J., Attatippaholkun, N., Kamkaew, M., Hou, S., Somasundaran, M., Wang, J.P., Finberg, R.W., Zhu, Q.K., et al. (2021). Unique structural solution from a V(H)3-30 antibody targeting the hemagglutinin stem of influenza A viruses. Nat Commun 12, 559.

Jardine, J., Julien, J.-P., Menis, S., Ota, T., Kalyuzhniy, O., McGuire, A., Sok, D., Huang, P.-S., MacPherson, S., Jones, M., et al. (2013). Rational HIV immunogen design to target specific germline B cell receptors. Science (New York, NY) 340, 711–716.

Jardine, J.G., Kulp, D.W., Havenar-Daughton, C., Sarkar, A., Briney, B., Sok, D., Sesterhenn, F., Ereño-Orbea, J., Kalyuzhniy, O., Deresa, I., et al. (2016). HIV-1 broadly neutralizing antibody precursor B cells revealed by germline-targeting immunogen. Science (New York, NY) 351, 1458–1463.

Jardine, J.G., Ota, T., Sok, D., Pauthner, M., Kulp, D.W., Kalyuzhniy, O., Skog, P.D., Thinnes, T.C., Bhullar, D., Briney, B., et al. (2015). HIV-1 VACCINES. Priming a broadly neutralizing antibody response to HIV-1 using a germline-targeting immunogen. Science (New York, NY) 349, 156–161.

Kallewaard, N.L., Corti, D., Collins, P.J., Neu, U., McAuliffe, J.M., Benjamin, E., Wachter-Rosati, L., Palmer-Hill, F.J., Yuan, A.Q., Walker, P.A., et al. (2016). Structure and Function Analysis of an Antibody Recognizing All Influenza A Subtypes. Cell 166, 596–608.

Kanekiyo, M., Joyce, M.G., Gillespie, R.A., Gallagher, J.R., Andrews, S.F., Yassine, H.M., Wheatley, A.K., Fisher, B.E., Ambrozak, D.R., Creanga, A., et al. (2019). Mosaic nanoparticle display of diverse influenza virus hemagglutinins elicits broad B cell responses. Nature immunology 20, 362–372.

Kepler, T.B. (2013). Reconstructing a B-cell clonal lineage. I. Statistical inference of unobserved ancestors. F1000Res 2, 103–103.

Koel, B.F., Burke, D.F., Bestebroer, T.M., van der Vliet, S., Zondag, G.C.M., Vervaet, G., Skepner, E., Lewis, N.S., Spronken, M.I.J., Russell, C.A., et al. (2013). Substitutions Near the Receptor Binding Site Determine Major Antigenic Change During Influenza Virus Evolution. Science (New York, NY) 342, 976–979.

Krause, J.C., Tsibane, T., Tumpey, T.M., Huffman, C.J., Basler, C.F., and Crowe, J.E., Jr. (2011). A broadly neutralizing human monoclonal antibody that recognizes a conserved, novel epitope on the globular head of the influenza H1N1 virus hemagglutinin. Journal of virology 85, 10905–10908.

Kuraoka, M., Schmidt, A.G., Nojima, T., Feng, F., Watanabe, A., Kitamura, D., Harrison, S.C., Kepler, T.B., and Kelsoe, G. (2016). Complex Antigens Drive Permissive Clonal Selection in Germinal Centers. Immunity 44, 542–552.

Lee, P.S., Ohshima, N., Stanfield, R.L., Yu, W., Iba, Y., Okuno, Y., Kurosawa, Y., and Wilson, I.A. (2014). Receptor mimicry by antibody F045–092 facilitates universal binding to the H3 subtype of influenza virus. Nature Communications 5, 3614.

Lee, P.S., Yoshida, R., Ekiert, D.C., Sakai, N., Suzuki, Y., Takada, A., and Wilson, I.A. (2012). Heterosubtypic antibody recognition of the influenza virus hemagglutinin receptor binding site enhanced by avidity. Proceedings of the National Academy of Sciences of the United States of America 109, 17040–17045.

Lingwood, D., McTamney, P.M., Yassine, H.M., Whittle, J.R.R., Guo, X., Boyington, J.C., Wei, C.-J., and Nabel, G.J. (2012). Structural and genetic basis for development of broadly neutralizing influenza antibodies. Nature 489, 566–570.

McCarthy, K.R., Watanabe, A., Kuraoka, M., Do, K.T., McGee, C.E., Sempowski, G.D., Kepler, T.B., Schmidt, A.G., Kelsoe, G., and Harrison, S.C. (2018). Memory B Cells that Cross-React with Group 1 and Group 2 Influenza A Viruses Are Abundant in Adult Human Repertoires. Immunity 48, 174-184.e179.

McCoy, A.J., Grosse-Kunstleve, R.W., Adams, P.D., Winn, M.D., Storoni, L.C., and Read, R.J. (2007). Phaser crystallographic software. Journal of Applied Crystallography 40, 658–674.

Pape, K.A., and Jenkins, M.K. (2018). Do Memory B Cells Form Secondary Germinal Centers? It Depends. Cold Spring Harb Perspect Biol 10.

Raymond, D.D., Bajic, G., Ferdman, J., Suphaphiphat, P., Settembre, E.C., Moody, M.A., Schmidt, A.G., and Harrison, S.C. (2018). Conserved epitope on influenza-virus hemagglutinin head defined by a vaccine-induced antibody. Proceedings of the National Academy of Sciences 115, 168.

Rohatgi, S., Ganju, P., and Sehgal, D. (2008). Systematic design and testing of nested (RT-)PCR primers for specific amplification of mouse rearranged/expressed immunoglobulin variable region genes from small number of B cells. J Immunol Methods 339, 205–219.

Sangesland, M., Ronsard, L., Kazer, S.W., Bals, J., Boyoglu-Barnum, S., Yousif, A.S., Barnes, R., Feldman, J., Quirindongo-Crespo, M., McTamney, P.M., et al. (2019). Germline-Encoded Affinity for Cognate Antigen Enables Vaccine Amplification of a Human Broadly Neutralizing Response against Influenza Virus. Immunity 51, 735-749.e738.

Sangesland, M., Yousif, A.S., Ronsard, L., Kazer, S.W., Zhu, A.L., Gatter, G.J., Hayward, M.R., Barnes, R.M., Quirindongo-Crespo, M., Rohrer, D., et al. (2020). A Single Human V(H)-gene Allows for a Broad-Spectrum Antibody Response Targeting Bacterial Lipopolysaccharides in the Blood. Cell reports 32, 108065.

Schmidt, Aaron G., Do Khoi T., McCarthy, Kevin R., Kepler, Thomas B., Liao, H.-X., Moody, M.A., Haynes, Barton F., and Harrison, Stephen C. (2015a). Immunogenic Stimulus for Germline Precursors of Antibodies that Engage the Influenza Hemagglutinin Receptor-Binding Site. Cell reports 13, 2842–2850.

Schmidt, Aaron G., Therkelsen Matthew D., Stewart, S., Kepler Thomas B., Liao, H.-X., Moody, M.A., Haynes Barton F., and Harrison, Stephen C. (2015b). Viral Receptor-Binding Site Antibodies with Diverse Germline Origins. Cell 161, 1026–1034.

Shlomchik, M.J. (2018). Do Memory B Cells Form Secondary Germinal Centers? Yes and No. Cold Spring Harb Perspect Biol 10.

Sui, J., Hwang, W.C., Perez, S., Wei, G., Aird, D., Chen, L.-m., Santelli, E., Stec, B., Cadwell, G., Ali, M., et al. (2009). Structural and functional bases for broad-spectrum neutralization of avian and human influenza A viruses. Nature structural & molecular biology 16, 265–273.

Tiller, T., Busse, C.E., and Wardemann, H. (2009). Cloning and expression of murine Ig genes from single B cells. J Immunol Methods 350, 183–193.

Viant, C., Weymar, G.H.J., Escolano, A., Chen, S., Hartweger, H., Cipolla, M., Gazumyan, A., and Nussenzweig, M.C. (2020). Antibody Affinity Shapes the Choice between Memory and Germinal Center B Cell Fates. Cell 183, 1298-1311.e1211.

Watanabe, A., McCarthy, K.R., Kuraoka, M., Schmidt, A.G., Adachi, Y., Onodera, T., Tonouchi, K., Caradonna, T.M., Bajic, G., Song, S., et al. (2019). Antibodies to a Conserved Influenza Head Interface Epitope Protect by an IgG Subtype-Dependent Mechanism. Cell 177, 1124-1135.e1116.

Whittle, J.R., Zhang, R., Khurana, S., King, L.R., Manischewitz, J., Golding, H., Dormitzer, P.R., Haynes, B.F., Walter, E.B., Moody, M.A., et al. (2011). Broadly neutralizing human antibody that recognizes the receptor-binding pocket of influenza virus hemagglutinin. Proceedings of the National Academy of Sciences of the United States of America 108, 14216–14221.

Whittle, J.R.R., Wheatley, A.K., Wu, L., Lingwood, D., Kanekiyo, M., Ma, S.S., Narpala, S.R., Yassine, H.M., Frank, G.M., Yewdell, J.W., et al. (2014). Flow Cytometry Reveals that H5N1 Vaccination Elicits Cross-Reactive Stem-Directed Antibodies from Multiple Ig Heavy-Chain Lineages. Journal of virology 88, 4047–4057.

Wrammert, J., Smith, K., Miller, J., Langley, W.A., Kokko, K., Larsen, C., Zheng, N.-Y., Mays, I., Garman, L., Helms, C., et al. (2008). Rapid cloning of high-affinity human monoclonal antibodies against influenza virus. Nature 453, 667–671.

Xu, R., Krause, J.C., McBride, R., Paulson, J.C., Crowe, J.E., Jr., and Wilson, I.A. (2013). A recurring motif for antibody recognition of the receptor-binding site of influenza hemagglutinin. Nat Struct Mol Biol 20, 363–370.

Yang, L., Caradonna, T.M., Schmidt, A.G., and Chakraborty, A.K. (2022). T helper cells select germinal center B cells stringently to promote the evolution of cross-reactive influenza antibodies. Submitted.

Yassine, H.M., Boyington, J.C., McTamney, P.M., Wei, C.-J., Kanekiyo, M., Kong, W.-P., Gallagher, J.R., Wang, L., Zhang, Y., Joyce, M.G., et al. (2015). Hemagglutinin-stem nanoparticles generate heterosubtypic influenza protection. Nature medicine 21, 1065–1070.

Yeh, C.-H., Nojima, T., Kuraoka, M., and Kelsoe, G. (2018). Germinal center entry not selection of B cells is controlled by peptide-MHCII complex density. Nature communications 9, 928–928.

Zost, S.J., Lee, J., Gumina, M.E., Parkhouse, K., Henry, C., Wu, N.C., Lee, C.-C.D., Wilson, I.A., Wilson, P.C., Bloom, J.D., et al. (2019). Identification of Antibodies Targeting the H3N2 Hemagglutinin Receptor Binding Site following Vaccination of Humans. Cell reports 29, 4460-4470.e4468.

