## Supplemental Information for "An epitope-enriched immunogen increases site targeting in germinal centers"





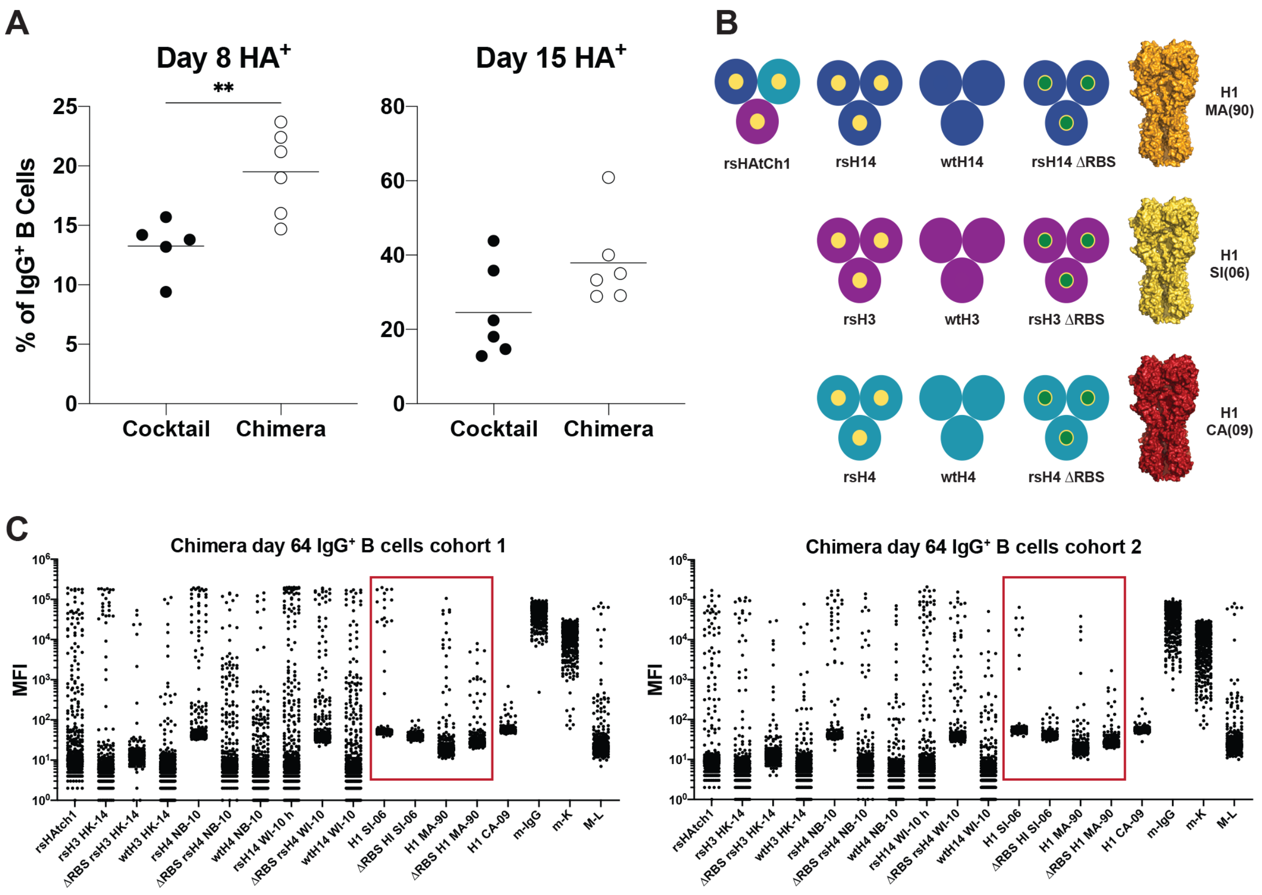


**Figure S1: Analysis of GC repertoires between cocktail and chimera immunization cohorts.** **(A)** Frequency of rHA+ B cells within the total GC B cell population. Combined results of GC analysis for cocktail and rsHAtCh-immunized cohorts at day 8 and day 15. **(B)** Cartoon diagram the recombinant antigen panel used for Nojima culture supernatant screening. **(C)** Mean fluorescent intensities (MFIs) for IgG+ culture supernatants being screened against a panel of recombinant HA antigens for two separate immunization cohorts (n = 3 each). Red box denotes H1 SI-06 and H1 MA-90 RBS-directed responses. For all data analysis, statistical significance determined using the Mann-Whitney U test, (**) denotes $p < 0.01$. **Related to Figure 1.**

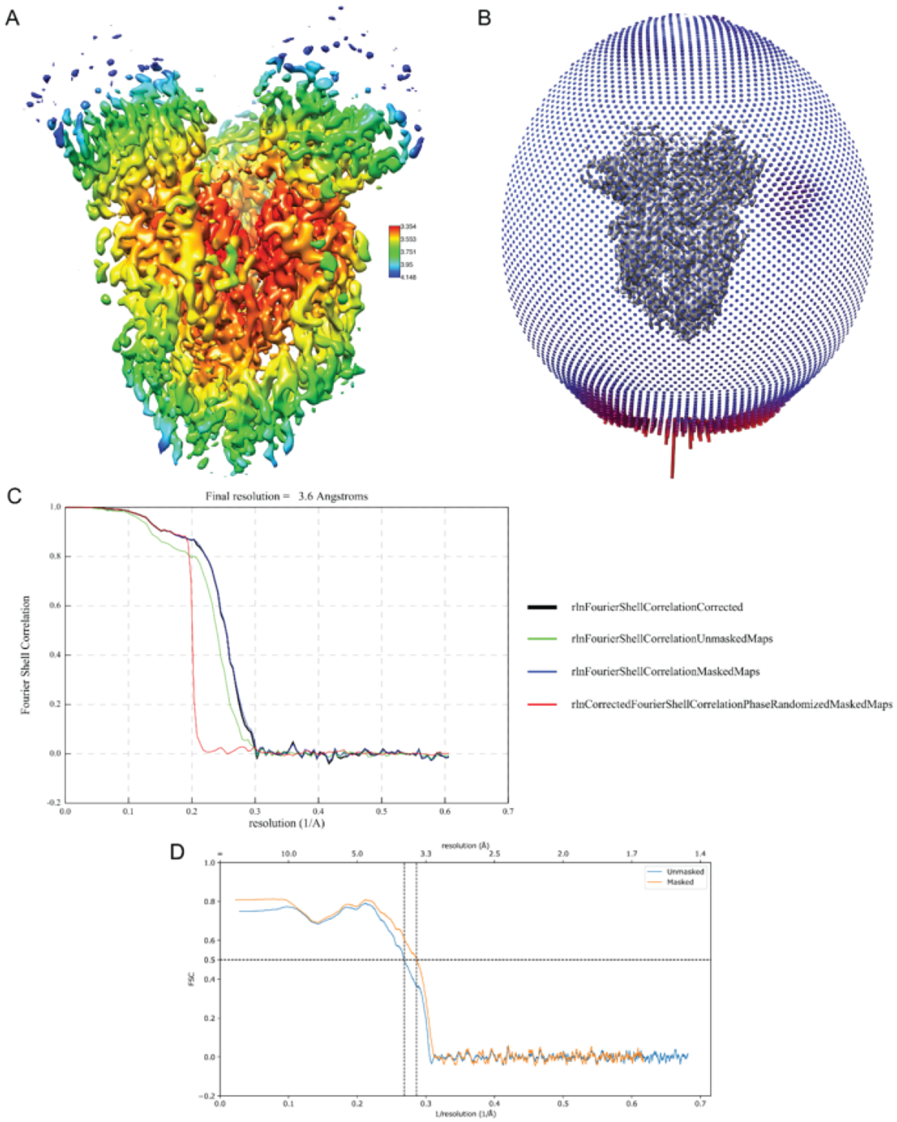


**Figure S2. Map and model quality for ab36 complex with H1 Solomon Islands 2006 HA.** A. The surface of the sharpened cryoEM density is colored based on resolution estimates generated by RELION. B. The angular distribution is shown from the final 3D refinement with RELION. C. FSC curves are shown for the final postprocessing in RELION. D. Map-to-model FSC curves are shown as generated by Phenix. **Related to Figure 6**

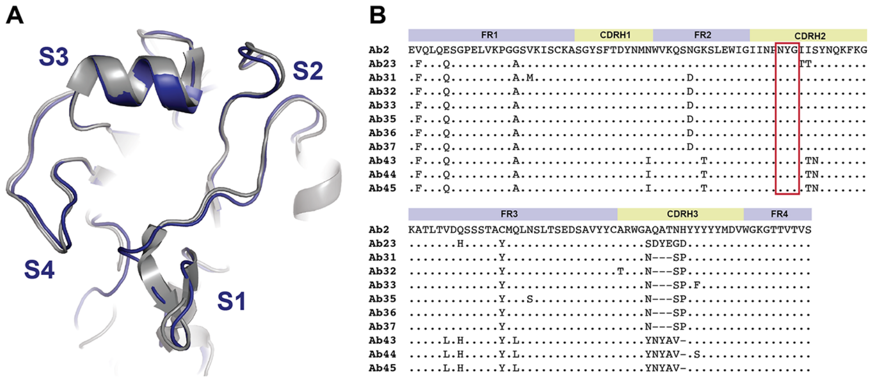


**Figure S3: $V_H$1-39-containing Abs engage structurally conserved RBS through a germline-encoded HCDR2 motif. (A)** Structural alignment of H1 SI-06 head (gray, PDB 4HKX) and rsH4 NB-10 (blue). Boundary segments of the grafted RBS epitope are shown in blue. **(B)** Sequence alignment of the eleven biochemically characterized $V_H$1-39-containing Abs, (.) denotes identity, (-) denotes a skip. The receptor-mimicking Asn-Tyr-Gly motif is marked with a red box.

**Table S1: Crystallography table for Ab27 Fab + rsH4 NB-10 head**

| | **rsH4 NB-10 head + Ab27 Fab** |
|---|---|
| **Wavelength (Å)** | 0.987 |
| **Resolution range** | 60.89 - 1.96 (2.03 - 1.96) |
| **Space group** | P $2_1$ $2_1$ $2_1$ |
| **Unit cell (Å)** | 63.215 71.034 226.613<br>90 90 90 |
| **Multiplicity** | 9.0 (6.2) |
| **Completeness (%)** | 98.20 (87.02) |
| **Mean I/sigma(I)** | 12.44 (1.01) |
| **Wilson B-factor** | 37.95 |
| **$R_{meas}$** | 0.187 (1.135) |
| **Reflections used in refinement** | 72952 (6378) |
| **Reflections used for $R_{free}$** | 1992 (175) |
| **$R_{work}$/$R_{free}$** | 0.2139 / 0.2273 |
| **Number of atoms** | 5953 |
| **Protein residues** | 709 |
| **RMS bonds (Å)/angles(°)** | 0.010 / 1.32 |
| **Ramachandran favored/allowed/outliers (%)** | 96.16 / 3.70 / 0.14 |
| **Clashscore** | 7.23 |
| **Average B-factor** | 44.12 |
| **Macromolecules** | 43.63 |
| **Ligands** | 150.71 |

Statistics for the highest-resolution shell are shown in parentheses. Related to **Fig. 6**

**Table S2.** Cryo-EM data collection and processing and model statistics

| Complex | H1 Solomon Island 2006·Ab36 Fab |
|---|---|
| Deposition | |
| **PDB ID** | |
| **EMDB ID** | |
| **Microscope** | FEI Titan Krios |
| **Voltage(kV)** | 300 |
| **Detector** | Gatan K3 |
| **Magnification (nominal)** | 105,000 |
| **Energy filter slit width** | 20 eV |
| **Calibrated pixel size (Å/pix)** | 0.825 |
| **Exposure rate (e- /pix/sec)** | 23.522 |
| **Frames per exposure** | 53 |
| **Total electron exposure ($e^-$/Å$^2$)** | 55.3 |
| **Exposure per frame ($e^-$/Å$^2$)** | 1.04 |
| **Defocus range (µm)** | -0.6, -2.0 |
| **Automation software** | Serial EM |
| **# of Micrographs used** | 5,274 |
| **Particles extracted** | 556,553 |
| **Total # of refined particles** | 124,426 |
| **Symmetry imposed** | C3 |
| **Map sharpening B-factor** | -149.3 |
| **Unmasked Resolution at 0.5/0.143 FSC (Å)** | 4.2/3.7 |
| **Masked resolution at 0.5/0.143 FSC (Å)** | 3.9/3.6 |
| Model refinement and validation | |
| Amino acids | 1584 |
| RMSD bond lengths (Å) | 0.013 |
| Angles (°) | 1.363 |
| Mean B-factors | 42.3 |
| Ramachandran Favored | 83.97 |
| Allowed | 16.03 |
| Outliers | 0.0 |
| Rotamer Outliers | 0.22 |
| Clash score | 28.13 |
| C-beta outliers | 0.0 |
| CaBLAM outliers | 6.84 |
| CC (mask) | 0.78 |
| MolProbity score | 2.61 |

Related to **Fig. 6**
